# Extracellular electrophysiology on clonal human β-cell spheroids

**DOI:** 10.1101/2024.04.02.587706

**Authors:** Emilie Puginier, Karen Leal Fischer, Julien Gaitan, Marie Lallouet, Pier-Arnaldo Scotti, Matthieu Raoux, Jochen Lang

**Author notes:** These authors contributed equally to this work and share first authorship (Emilie Puginier, Karen Leal Fischer).

## Abstract

Pancreatic islets are important in nutrient homeostasis and improved cellular models of clonal origin may very useful especially in view of relatively scarce primary material. Close 3D contact and coupling between β-cells are a hallmark of physiological function improving signal/noise ratios. Extracellular electrophysiology using micro-electrode arrays (MEA) is technically far more accessible than single cell patch clamp, enables dynamic monitoring of electrical activity in 3D organoids and recorded multicellular slow potentials (SP) provide unbiased insight in cell-cell coupling. We have therefore asked whether 3D spheroids enhance clonal β-cell function such as electrical activity and hormone secretion using human EndoC-βH1, EndoC-βH5 and rodent INS-1 cells. EndoC-βH1 spheroids exhibited increased signals in terms of SP frequency and especially amplitude as compared to monolayers and even single cell action potentials (AP) were quantifiable. Enhanced electrical signature in spheroids was accompanied by an increase in the glucose stimulated insulin secretion index. EndoC-βH5 monolayers and spheroids gave electrophysiological profiles similar to EndoC-βH1, except for a higher electrical activity at 3 mM glucose, and exhibited moreover a biphasic profile. Again, physiological concentrations of GLP-1 increased AP frequency. Spheroids also exhibited a higher secretion index. INS-1 cells did not form stable spheroids, but overexpression of connexin 36, required for cell-cell coupling, increased glucose responsiveness, dampened basal activity and consequently augmented the stimulation index. In conclusion, spheroid formation enhances physiological function of the human clonal β-cell lines and these models may provide surrogates for primary islets in extracellular electrophysiology.

## 1 Introduction

Pancreatic islets are important in nutrient homeostasis and their dysfunction leads to a major metabolic disease, diabetes (1, 2). Glucose metabolism in β-cells leads to membrane depolarisation and calcium influx that triggers insulin secretion (3). Studies on primary islet cells are hampered by the relative scarceness of native material, especially those of human origin, which moreover differ in several important aspects from rodent islet cells (3)and exhibit a high degree of variability (4). Consequently, clonal β-cell lines provide useful models and this approach has been considerably improved by the establishment of a human β-cell line, EndoC-βH1 cells and their derivatives (5-9).

A number of parameters such as ultrastructure, gene expression, survival and secretion has been investigated in EndoC-β cells, however, electrical activity has only been addressed in EndoC-βH1 and -βH2 and only in single cell analysis in monolayers (7, 10), which are not fully informative of their properties in 3D conformation or in islets. Thus, important parameters of function remain largely unknown. Extracellular electrophysiology such as micro-electrode arrays (MEA) offers a direct and non-biased approach on whole islet characteristics. Recorded Slow potentials (SP) (11, 12) are multicellular events strictly depending on gap junction coupling by Cx36 in islet cells and their amplitude reflects the degree of coupling (10, 11, 13, 14). Furthermore, the electrical activity determined by MEAs is closely correlated to secretion, and its glucose concentration dependency enables small increases in glucose to be distinguished in both human and murine islets (11, 13-15).

Native islets are organoids and considerable effort has been spent to assemble clonal β-cells in 3D aggregates or spheroids. Such an assembly should increase contacts between β-cells and coupling between β-cells mediated by connexin 36 (Cx36) are of importance for physiological responses (16, 17). Cx36 form gap junctional channels that provide electrical coupling between β-cells and influencing synchronisation (18-20). Cx36 mediated coupling not only entrains cells upon arrival of a stimulus, but also dampens hyperactive cells and thus reduce spontaneous activity (21-23) which results in an improved signal/noise ratio. In contrast to primary β-cells, connexin 36 expression is generally low in β-cell lines (24).

A number of approaches have been used to generate spheroids from clonal β-cells (25, 26) such as specific media and plating in proprietary wells or microgravity (hanging drop) for human EndoC-β cells (27-29) or rodent β-cell lines (30-33). Since recording spheroids by MEAs requires electrical contact, this precludes certain methods of spheroid formation such as coculture with endothelial cells (34) or encapsulation (35). However, using electrical activity as read-out for activity and coupling offers certain advantages as compared to fluorescent approaches such as absence of bleaching or destructive analytical methods. It is also easier to miniaturize and to multiplex, and signals can even be analysed online (13, 36).

Using 3D spheroids and MEAs we have now determined effective coupling in EndoC-βH1 and especially EndoC-βH5 cells. Our data indicate that stimulus-dependent coupling is considerably enhanced in 3D spheroids thus providing a base for their improved activity. In contrast, a widely used rat clonal β-cell line, INS-1, did not form stable spheroids. Nevertheless, responsiveness and activity were considerably enhanced by Cx36 overexpression. Therefore, both models may provide paradigms to explore mechanisms and test drugs that rely on physiological β-cell coupling.

## 2. Materials and Methods

### 2.1. Materials

EndoC-βH1 cells (5) were kindly provided by Human Cell Design (Toulouse, France), EndoC-βH5 cells were purchased from Human Cell Design (Toulouse, France) and cultured according the manufacturer’s instructions. INS832/13 cells were cultured as described previously (37, 38). IBMX, forskolin and glibenclamide were purchased from Sigma, GLP-1 from Bachem (Bubendorf, Switzerland). The following primary antibodies were used (all at 1:100 dilution in immunofluorescence, 1:1000 in immunoblots): CX36 mouse anti-human (Invitrogen, clone 1E5H5), rabbit recombinant ANTI-FLAG M2 antibody (Invitrogen 710662), guinea pig anti-bovine insulin (Linco, St. Charles, MO, USA), monoclonal anti-insulin (Sigma, clone K36AC10), monoclonal anti-glucagon (Sigma, clone K79bB10), polyclonal goat anti-somatostatin (Santa Cruz, sc-7819), monoclonal anti-GFP, monoclonal anti-SNAP-25 (SP12, Sternberger Monoclonals) or monoclonal anti-VAMP2 (Synaptic Systems, Göttingen, Germany). The following secondary antibodies were used: anti-mouse or anti-rabbit HRP (dilution1/2000;

GE Healthcare); anti-mouse or anti-rabbit alexa568 (dilution 1/300; Invitrogen A11012 and A11031), anti-goat TMR, donkey anti-guinea pig (Jackson Laboratories, Bar Harbour, ME, USA). Note that two other primary polyclonal antibodies did not provide any reliable signal in islets or brain for Cx36 (Invitrogen 701194 and 516300). pLenti-C-Myc-DDK (RC210158L1; carrying the ORF of human CX36; GJD2; NM_020660) was obtained from Origene (Rockville, Md, USA).

### 2.2. Cell Culture and spheroid formation

EndoC-βH1 and EndoC-βH5 cells (5) were cultured according to the manufacturers protocol in OPTIβ1 (Human Cell Design, Toulouse, France). INS-1E and INS-813 cells were cultured as described previously (39, 40) and primary mouse islets (male C57BL/6, age 16-24 weeks) were prepared and cultured as published (12, 14, 41). Spheroids were formed in complete medium either by hanging drop for 5 days in 30 µl containing 500 islet cells or using a commercial plate (Sphericalplate 5D, Kugelmeiers; Erlenbach, Switzerland) with indicated cell concentrations.

Physical stability was tested by 10 times pipetting trough 200 µl tips and visual inspection with a microscope. Spheroids were considered as stable if no disaggregation was observed. Spheroid dimensions were determined on microscopic images using ImageJ v1.53.

### 2.2. Viral transduction and quantitative PCR

Lentiviral vector production was done by Vect’UB of the Bordeaux University. Lentiviral vector was produced by transient transfection of 293T cells according to standard protocols. In brief, subconfluent 293T cells were cotransfected with lentiviral genome (psPAX2) (42), with an envelope coding plasmid (pMD2G-VSVG) and with vector constructs. Viral titers of pLV lentivectors were determined by transducing 293T cells with serial dilutions of viral supernatant and EGFP expression was quantified 5 days later by flow cytometry analysis. For transduction of INS-1 cells, 750.000 cells were incubated in 500 µl of RPMI and 5 MOI of corresponding viral particles overnight, washed and placed in complete RPMI medium for 5 days prior to plating.

### 2.3. Secretion assays and immunocytochemistry

Static secretion assays were performed as described (43) using Krebs-Ringer bicarbonate HEPES buffer (KRBH, concentrations in mM, 135 NaCl, 3.6 KCl, 5 NaHCO3, 0.5 NaH2PO4, 0.5 MgCl2, 1.5 CaCl2, 10 HEPES, 0.1% w/v BSA, pH 7.4) and commercial ELISAs (Mercodia, Uppsala, Sweden). Immunocytochemistry was performed as described (44) and images acquired with a CAMSCOP CMOS camera (SCOP-Pro, Ballancourt, France) linked to an inverted fluorescent microscope (TE 200, Nikon; Champigny, France).

### 2.4. Electrophysiology

MEA recordings (60Pedot-MEA200/30iR-Au-gr, Ø30 µm, 200 µm inter-electrode distance; MCS, Tübingen, Germany) were performed at 37°C in solutions containing (in mM) NaCl 135, KCl 4.8, MgCl2 1.2, CaCl2 1.2 or 2.5 in the case of INS cells HEPES 10 and glucose as indicated (pH 7.4 adjusted with NaOH) (11, 14, 15, 37, 45). MEAs were coated with Matrigel (2% v/v) (BD Biosciences, San Diego, CA) prior to seeding of cells, spheroids or islets. Electrodes with noise levels >30 µV peak-to-peak were regarded as artefacts, connected to the ground and not analysed. Extracellular field potentials were acquired at 10 kHz, amplified and band-pass filtered at 0.1-3000 Hz with a USB-MEA60-Inv-System-E amplifier (MCS; gain: 1200) or a MEA1060-Inv-BC-Standard amplifier (MCS; gain: 1100) both controlled by MC_Rack software (v4.6.2, MCS) (12, 14).

### 2.5. Quantitative real-time PCR (qPCR)

Quantitative PCR was performed as described previously (38). *YHWAZ* (Tyrosine 3-Monooxygenase/Tryptophan 5-Monooxygenase Activation Protein Zeta) and *GAPDH* were used as reference genes. *FAP* (Fibroblast activation protein, alpha), *IRX2* (iroquois homeobox 2) and *GCG* (glucagon) were used as marker genes for β-cells (46). Details and primers used are given in Supplemental Table 1.

### 2.6. Statistics

Data are presented as means and SD except for mean traces where SEMs were given to enhance readability. Gaussian distributions were tested by Shapiro-Wilk test and one-way ANOVA with Tukey post hoc or nonparametric Dunn tests were used; n corresponds to the number of electrodes recorded (from 3 distinct experiments).

## 3. Results

### 3.1. Human EndoC-βH1 cell spheroids

Spheroids of human EndoC-βH1 cells were generated using micro-structured plastic wells for culture and the assembly and growth properties were tested first. As given in Fig. 1A.i, EndoC-βH1 cells formed round and regular spheroids with a rather homogenous staining for insulin. Different spheroid sizes were monitored according to initial cell number seeding and to incubation time. Based on our experience with the culture of islets, which are native spheroids, we opted for an intermediate diameter of 100 µm (Fig. 1 A.ii) and used the corresponding protocol for all further experiments. These spheroids were mechanical stable after repetitive pipetting and conserved their spheroid form during culture on microelectrode arrays (Fig. 1 A.iii). Pseudoislets from primary mouse islet cells were prepared by the hanging drop method as this approach uses smaller quantities of cells (Fig. 1 B).

**Figure 1:**
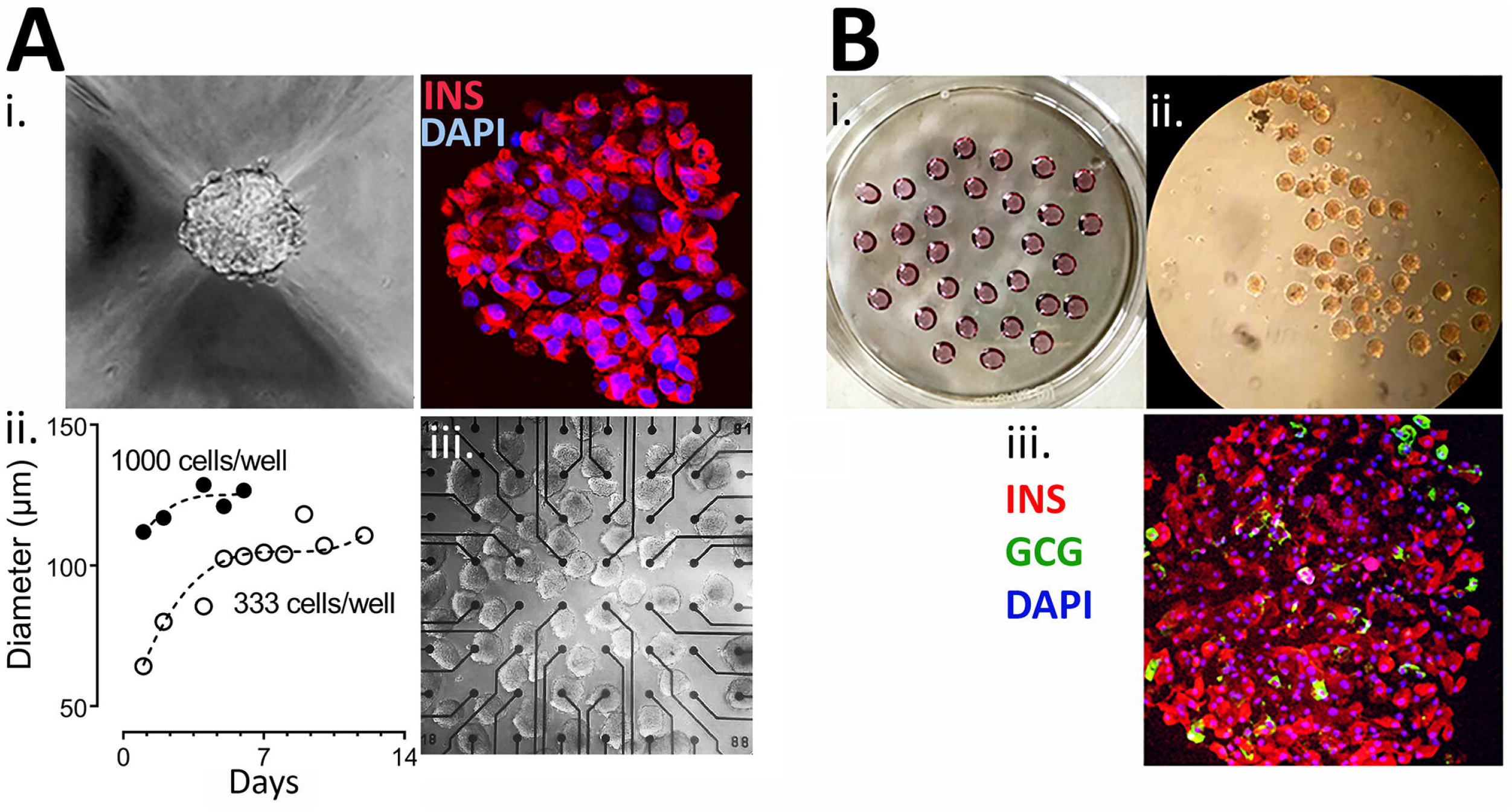
Generation of Spheroids. A: Generation of EndoC-βH1 3D spheroids. i, image of spheroid formed and staining for insulin (red) and with DAPI (blue); ii, time course of spheroid formation and size at different cell numbers; iii, EndoC-βH1 3D spheroids on a micro-electrode array. B, formation of islet spheroids. i petri with hanging drops; ii, 3D islets formed; iii, staining for insulin, glucagon and with DAPI.x

We next examined their electrical activity by comparing 2D monolayer versus spheroids of EndoC-βH1 on microelectrode arrays. A hallmark of physiological β-cell behaviour (16, 47), the so-called slow potentials (SP) were detected and reflect activity of electrically coupled cells (11). In addition, the amplitude of SPs is related to the degree of electrical coupling and synchronization between the cells within an islet (11, 14). Raising glucose from 3 to 11 mM significantly increased SP frequency and amplitude in 2D culture and in spheroids and the mean effect was significantly more pronounced in the latter (Fig. 2A, B). Interestingly EndoC-βH1 spheroids had a higher basal activity (3 mM glucose) and thus fold increase between G3 and G11 in terms of mean frequency was more pronounced in monolayers (5.7 fold vs 1.8 fold in spheroids), but stronger in terms of amplitude in spheroids (1.6 in spheroids vs 1.2 fold in monolayers). The further addition of the incretin GLP-1 at the physiological concentration of 50 pM induced a slight further increase in both cases which did not reach significance. Increasing cellular cAMP levels by the direct adenylate cyclase activator forskolin and the phosphodiesterase inhibitor IBMX, in the presence of 11 mM glucose, significantly increased frequencies as compared to 11 mM glucose alone in 2D culture and spheroids. Similar effects of forskolin/IBMX were observed for amplitudes, and again the effects were considerably more pronounced in spheroids as compared to 2D cultures (Fig. 2A, B).

**Figure 2:**
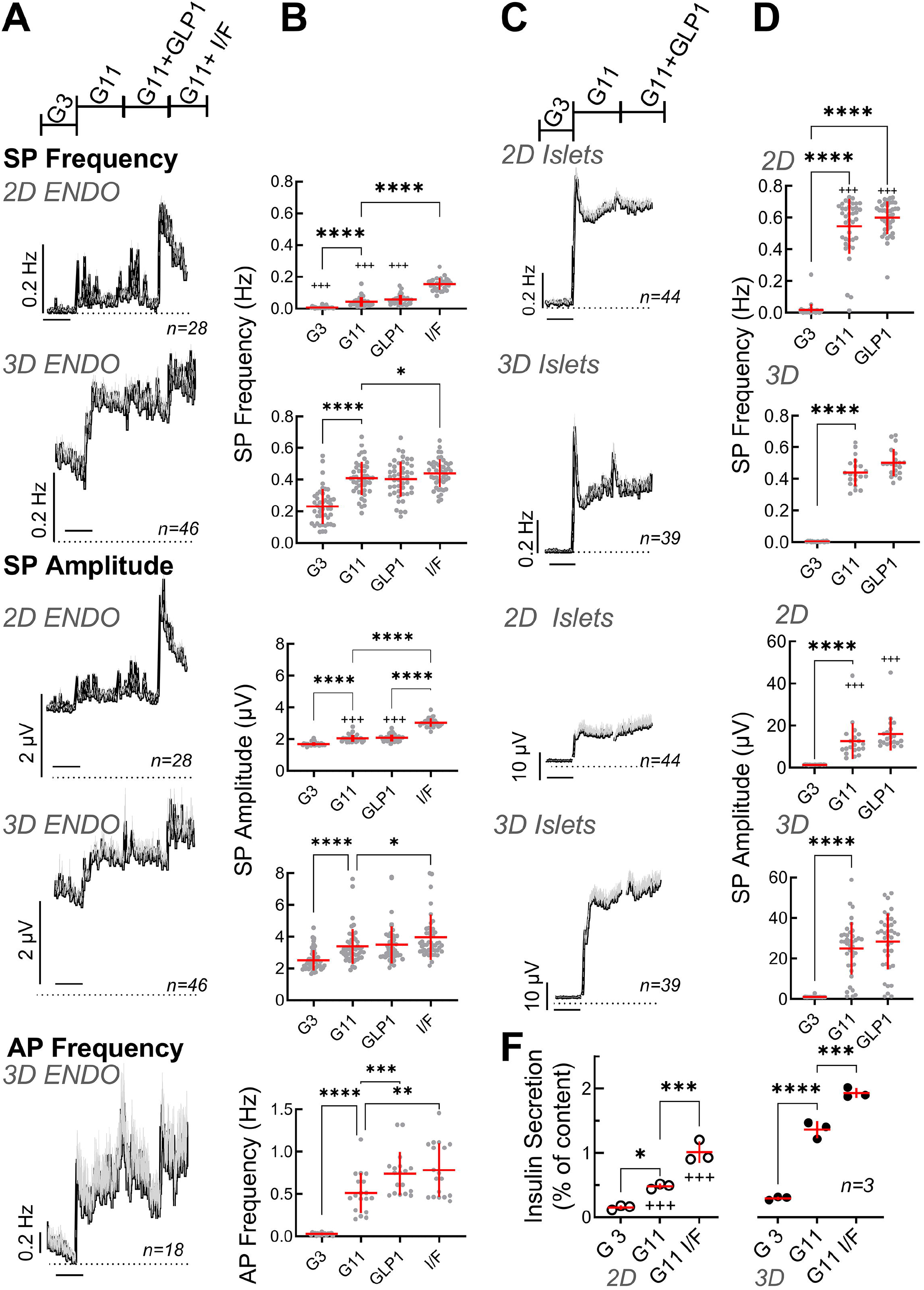
Functional characterisation of spheroids from EndoC-βH1 cells or primary mouse islets. A: Recording of monolayer (2D) or spheroids (3D) of EndoC-βH1 cells seeded on micro-electrode arrays and exposed to Glucose (3 mM, G3; 11 mM, G11), GLP-1 (50 pM) in the presence of 11 mM glucose (GLP1) or IBMX (100 µM) and Forskolin (1 µM) in the presence of 11 mM glucose (I/F). Mean traces of slow potential (SP) frequency and amplitudes as well as action potential (AP) frequency are given; mean, black, SEM grey. Time bars equal 20 min (in all traces). B: Statistical evaluations of the curves of A (mean values). C: Recording of monolayer (2D) or reassembled spheroids (3D) of primary mouse seeded on micro-electrode arrays. Abbreviations for conditions and statistical tests as in A. D: Statistical evaluations of curves of C (mean values). E: Action potential frequencies in spheroids and statistical analysis. F: Insulin secretion (static incubations) of monolayer (2D) or spheroids (3D) of EndoC-βH1 cells during 1h incubation, abbreviations as in A. ANOVA and Tukey posthoc test; *, 2p<0.05; ** 2p<0.01, *** 2p<0.001, ****2p<0.0001; 2D vs 3D, ^+++^ 2p<0.001. n, given in corresponding panels.

In contrast to the very robust SP signals, single cell action potentials are more difficult to detect in conventional MEAs and their amplitudes cannot be reliably determined (45). Moreover, as only the frequency but not the amplitude of APs varies with glucose stimulation (12), we only analysed their frequency. In 2D cultures we were not able to identify APs with certainty, probably due to their extremely low amplitude. In contrast, recordings of spheroids clearly showed APs which increased with a raise in glucose concentration and significant effects of GLP-1 as well as effects of IBMX/forskolin were observed as compared to elevated glucose alone.

As a comparison to EndoC-βH1 cells we examined also mouse islet cells, either as dispersed single cell 2D culture or after reaggregation in spheroids (Fig. 2 C, D). In both cases, large effects were present in terms of frequency and amplitude when increasing glucose from 3 to 11 mM and the glucose-induced increase in frequency was clearly biphasic described previously (14). A small transient effect effect was observed for GLP-1 (50 pM) in the presence of 11 mM glucose. As already observed for the EndoC-βH1 cell line, the effect of glucose on primary islet cells was more pronounced in terms of frequency in monolayers (139 vs. 89 fold) and stronger in terms of amplitudes in spheroids (25 vs. 8.9 fold).

Next, we measured insulin secretion from 2D cultures and 3D EndoC-βH1 spheroids (Fig. 2 E). Glucose-induced stimulation was clearly apparent as well as further potentiation by IBMX/forskolin. Similar to electrical activity, basal release and stimulated insulin secretion was more pronounced in spheroids as compared to 2D cultures. Thus, the stimulation index increased from 2.6 in monolayers to 4.5 in spheroids (see Supplemental Figure 1). For comparison, we also measured static secretion in primary mouse islets were a rise in glucose from 0 to 3 and 11 mM increased insulin secretion from 0.05+0.02 to 0.07+0.02 and 1.15+0.19 (percent of content, n=5; stimulation index SI 16.4).

### 3.2. Human EndoC-βH5 cell spheroids

We subsequently tested an optimized EndoC-βH version (Fig. 3), the EndoC-βH5 cells, known for their improved function (9). EndoC-βH5 easily formed stable spheroids (Supplemental Fig. 2) and a mean diameter of 120 µm was used. Immunofluorescence demonstrated the expected presence of insulin as well as the SNARE proteins VAMP2 and SNAP-25. As reported previously (9), EndoC-βH5 also stained for glucagon and somatostatin in spheroids (Supplemental Fig. 2) as did monolayers (data not shown). The presence of glucagon was also detected by qPCR (Supplemental Figure 3) although other markers for α-cells, such as FAP and IRX were not detectable. Note also that we could not detect any glucagon in static secretion assays (data not shown). Spheroid formation in EndoC-βH5 did not change the expression of glucagon or connexin 36 (GJD2) and a small decrease in the expression of preproinsulin was apparent.

**Figure 3:**
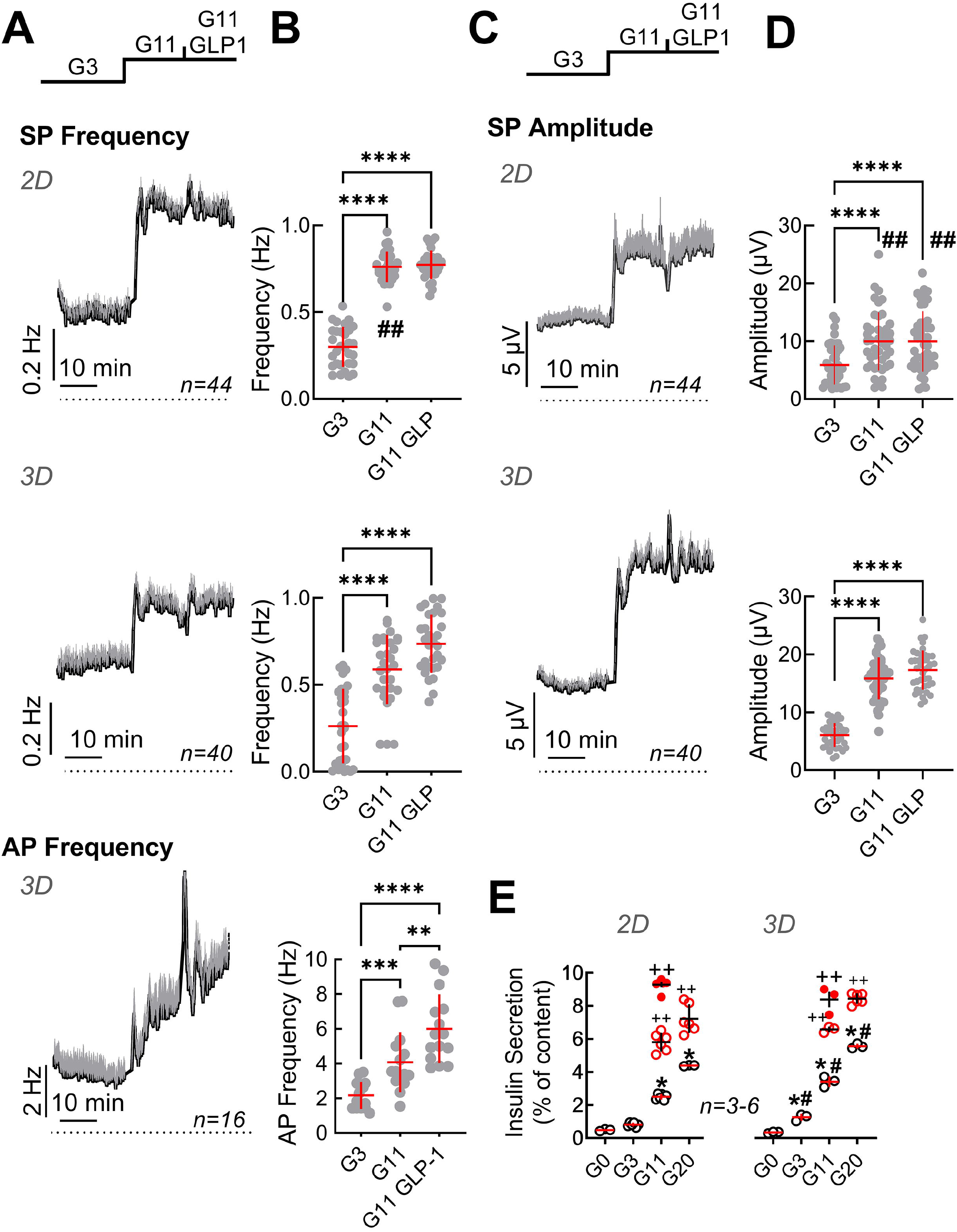
Functional characterisation of monolayers and spheroids from EndoC-βH5 cells. Recording of monolayer (2D) or spheroids (3D) of EndoC-βH5 cells seeded on micro-electrode arrays and exposed to Glucose (3 mM, G3; 11 mM, G11) or GLP-1 (50 pM) in the presence of 11 mM glucose (G11 GLP1). A: Mean traces of slow potential (SP) frequency as well as action potential (AP) frequency bare given; mean, black, SEM grey. Time bars equal 10 min (in all traces). B: Statistical evaluations of mean frequencies of data given in A. C: Mean traces of slow potential (SP) amplitudes; mean, black, SEM grey. D: Statistical evaluations of data given in C. E: Insulin secretion (static incubations) of monolayer (2D) or spheroids (3D) of EndoC-βH5 cells during 1h incubation, abbreviations as in A. Open and filled red circles, IBMX (0.1 mM)/Forskolin (1 µM) or GLP-1 (50 pM) in the presence of indicated concentrations of glucose.; ANOVA and Tukey posthoc test; B, D, E: *, 2p<0.05; **, 2p <0.01; ***, 2p <0.001, ****, 2p <0.0001; Comparison 2D and 3D: #, 2p<0.05, ##, 2p<0.01; insulin secretion (E), ++, 2p <0.01 as compared to the absence of GLP-1 or IBMX /Forskolin; n, given in corresponding panels.

Slow potentials were more pronounced in EndoC-βH5 as compared to EndoC-βH1 in terms of frequencies and amplitudes for both, monolayers and spheroids (Fig. 3). Similarly, action potential frequencies were higher and an effect of GLP-1 was measurable. However, an increased basal activity (at G3) was also evident in both, monolayers and spheroids, as compared to EndoC-βH1 and thus the fold increases of mean activities in terms of amplitude and frequency between G3 and G11 were comparable between the two cell lines. As observed for the EndoC-βH1 cells, spheroid formation reduced the fold increase in frequency from 3 to 11 mM glucose (2.2 fold in monolayers vs 1.8 fold in spheroids) and augmented the fold-increase in amplitudes (2.5 fold in monolayers vs 3.5 fold in spheroids). The observed temporal development of slow potentials in EndoC-βH5 cells may also suggest the presence of a first peak indicating biphasic behaviour (Fig. 3) although this was clearly less pronounced as compared to primary mouse β-cells (see Fig. 2 C). Similar to EndoC-βH1 cells, spheroid formation enhanced insulin secretion in EndoC-βH5 (Fig. 3 E) and the cells responded well to additional GLP-1 or IBMX/Forskolin. The stimulation index of insulin secretion increased from 5-(monolayers) to 10-fold (spheroids) when compared between 0 and 20 mM glucose, but between 3 and 11 mM glucose the stimulation index was around 3 fold and thus comparable to EndoC-βH1. We also noted that spheroids, in difference to monolayers, increased hormone secretion already at 3 mM glucose as compared to the absence of glucose.

### 3.3. Rat INS-1 cell spheroids

We also tried to generate spheroids from another frequently used cell line, i.e. rat insulinoma derived INS-1 cells (37, 48-50). However, these spheroids proved to be unstable to repetitive pipetting and even when handled with considerable care, spheroids rapidly disaggregated when cultured on MEAs precluding their use in 3D conformation (data not shown). Moreover, in 2D cultures a considerable number of cells or cell clusters did not respond in terms of measurable electrical activity upon increases of glucose. We therefore tested whether an increase in the expression of CX36, required for intercellular coupling and participating in adhesion (51), may improve their electrical responses.

To this end INS-1 cells were transduced with viral particles encoding either GFP as a control or human connexin 36 (CX36). Immunoblot analysis of non-transduced cells and cells transduced with GFP or CX36 revealed expression of GFP or of CX36 only in the correspondingly transduced cells as bands appearing at approximately 25 kDa (GFP) or around 36 kDa as well as 110 kDa trimers (CX36) upon co-staining with GFP- and CX36 antibodies (Fig. 4A). Human connexin was expressed intracellularly and also fine rims could be observed compatible with location at the plasma membrane (Fig. 4 B). In contrast, incubation with the anti-connexin antibody did not reveal any staining in GFP-transduced cells (Fig. 4 B).

**Figure 4:**
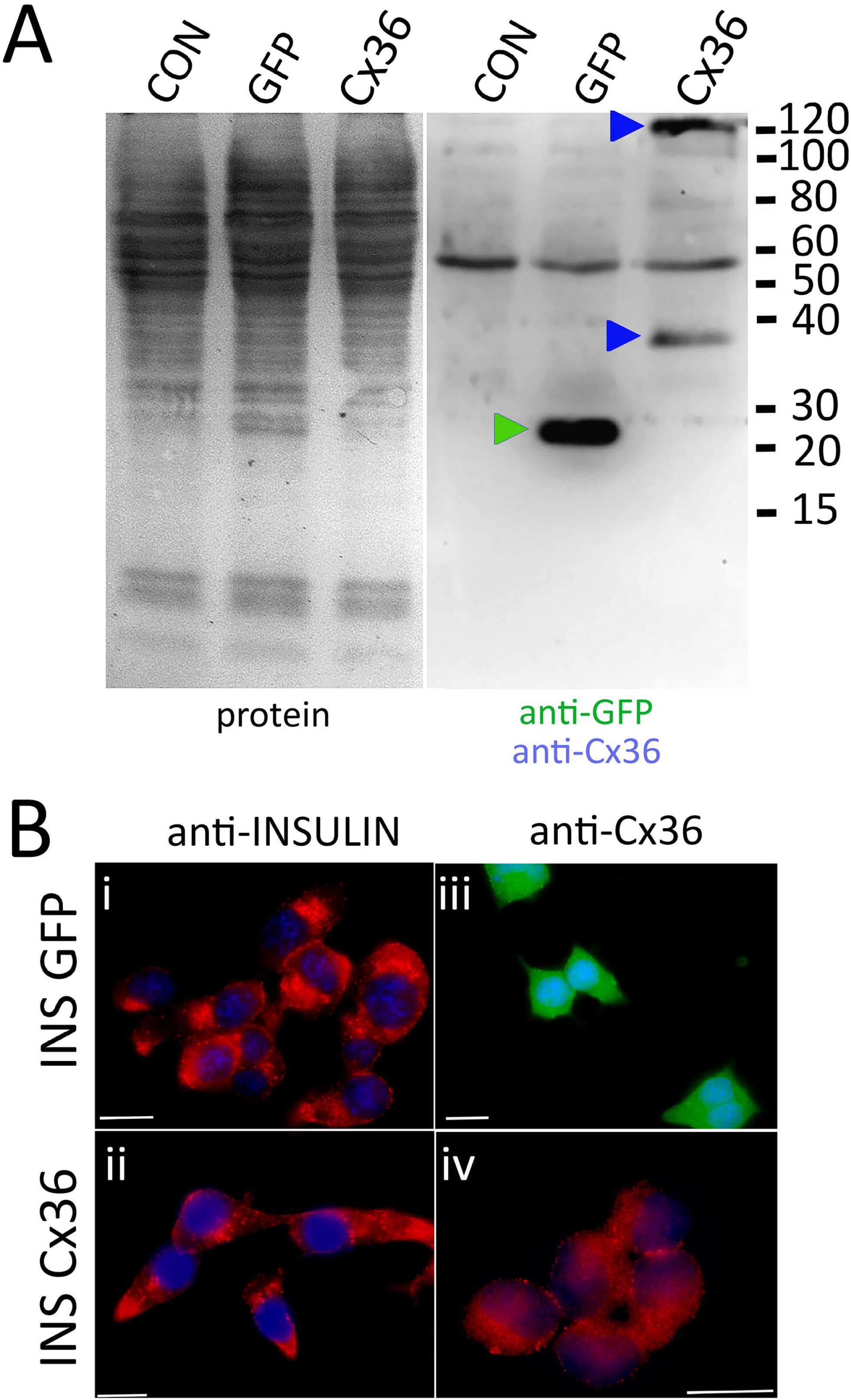
CX36 expression in transduced INS-1 cells. A, immunoblot of non-transduced cells (CON) or cells transduced with either eGFP (GFP) or connexin-36 tagged with a myc-epitope (Cx 36). Left panel, protein transfer; right panel, corresponding blot co-incubated with anti-eGFP and anti-myc. Molecular weight markers are given in kDa, specifically labelled bands are indicated by correspondingly coloured triangles. B, Clonal INS-1 images of INS-1 β-cells were transduced with viral particles encoding either eGFP (INS GFP, i and iii) or with Cx36 (INS Cx 36, ii and iv) and stained for insulin (i, ii) or for Cx36 (iii, iv). GFP expression was detected directly. Bars, 10 µm.

We subsequently compared the electrical responses in terms of SP frequency and amplitude of GFP- and of Cx36-transduced cells in response to 3 or 11 mM glucose and a mix of stimulatory drugs (glibenclamide 200 nM, Bay K8644 1 µM, forskolin 1 µM) in the presence of 11 mM glucose (Fig. 5). We first observed a considerable difference in their reactivity in terms of electrodes covered with cells which recorded changes in electrical activity (Fig. 5B). Whereas in GFP-transduced cells only a minority of cells responded to an increase in glucose, more than a half were active using connexin-36 transduced cells. In fact, most of the GFP-transduced cells did not respond to glucose or glucose in the presence of stimulatory drugs (glibenclamide, Bay K8644, forskolin) in line with observations from cultures of native INS-1 cells (data not shown). We subsequently analysed in detail the recordings from those electrodes covered with cells that responded at least to an increase in glucose from 3 to 11 mM (Fig. 5, C-F). In both, GFP- or Cx36-transduced cells, the change from complete culture medium to 3 mM glucose reduced activity in terms of frequency and amplitudes. Note that complete culture medium contains 11 mM glucose and amino acids, the latter being known to enhance glucose effects (50). Interestingly, at low glucose (3 mM), SP frequency was significantly lower in Cx36-transduced cells as compared to GFP-transduced cells (Fig. 5 D). Subsequent change from 3 to 11 mM glucose increased slightly but not significantly frequency and amplitude in GFP-transduced cells whereas a significant effect was observed in CX36-transduced cells. Further exposure to stimulatory drugs significantly increased responses in CX36-transduced cells, whereas only amplitude but not frequency was enhanced in GFP-transduced cells.

**Figure 5:**
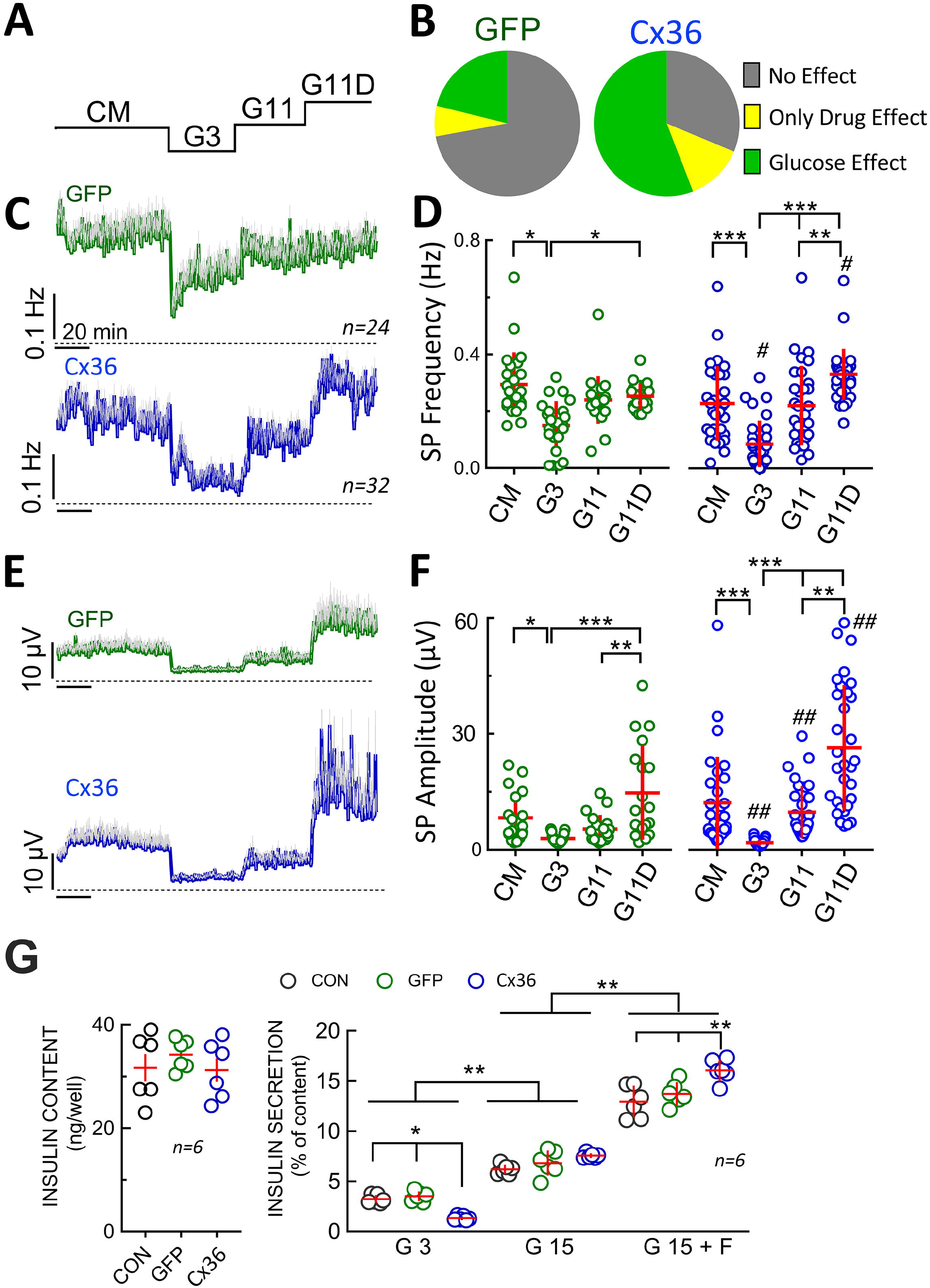
Electrophysiological analysis and insulin secretion of GFP or CX36 expressing transduced INS-1 cells. A, Scheme of static incubation of INS-1 cells with culture medium (CM), 3 or 11 mM glucose (G3, G11) or 11 mM glucose in the presence of drugs (by glibenclamide 200 nM, Bay K8644 1 µM, forskolin 1 µM). B, relative responsivity of GFP- or Cx36 (Cx36) transduced cells expressed as absence of effect, stimulation by 11 mM glucose (versus 3 mM) or only stimulated by drugs (no effect of G11 alone; increase versus G3 by glibenclamide 200 nM, Bay K8644, forskolin 1 µM). Note that glucose-sensitive cells were always also drug sensitive. For further analysis (C-F) only those electrodes covered by cells were analysed where an increase in glucose increased electrical activity. C, Mean SP frequencies (+SEM) in GFP- or Cx36 transduced cells. D, statistics of C. E, Mean SP amplitudes (+SEM) in GFP- or Cx36 transduced cells. F, statistics of E. G, Insulin content and insulin secretion from non-transduced cells (CON) or GFP-(GFP) or Cx36 (Cx36) transduced cells incubated at 3 mM glucose (G3), 15 mM glucose (G15) or 15 mM glucose and 1 µM forskolin (G15 F). Statistics: Tukey or Dunn post-hoc tests; *, 2p<0.05; **, 2p <0.01; ***, 2p <0.001; Comparison 2D and 3D: #, 2p<0.05, ##, 2p<0.01; n, given in corresponding panels.

Finally, we determined insulin secretion in non-transduced and GFP- or Cx36-transduced cells. Under all three conditions (non-transduced, GFP-transduced or CX36 transduced cells) insulin content did not vary. Clearly, Cx36 expression reduced basal secretion (at 3 mM glucose) in Cx36 transduced cells as compared to the two other conditions. In all three cell types, an increase in glucose stimulated secretion was observed. The stimulation index (15 mM vs 3 mM glucose) amounted to 1,9 in non-transduced and GFP transduced cells, but increased to 5,7 in Cx36 transduced cells. Although Cx36-transduced cells secreted 20% more insulin than GFP-transduced cells at 15 mM glucose, the increase in GSIS was mainly due to an approximately 60% reduction in basal secretion at 3 mM glucose in Cx36-transduced cells. Forskolin in the presence of 15 mM glucose further enhanced insulin secretion and again to a greater degree extent in Cx36-transduced cells as compared to GFP-transduced or non-transduced cells.

## 4. Discussion

Our results indicate that spheroids of human EndoC-βH1 and -βH5 cells exhibit more pronounced signals in extracellular physiology than monolayers. Spheroids have a relatively lower frequency and higher amplitude of slow potentials indicating a higher degree of cell-cell coupling. This is accompanied by a considerable improvement in the glucose-stimulated secretion index. The lack of stable spheroid formation in rat clonal INS-1 β-cells may be compensated by the enhanced expression of connexin 36, which also improves GSIS index mainly by lowering basal electrical activity and ensuing secretion.

3D spheroids of human EndoC-βH1, -βH3 or -βH5 were generated previously using microgravity (“hanging drop”), low attachment plates or co-culture on human umbilical vein or islet-derived endothelial cells (9, 27-29, 52). These spheroids exhibited a GSIS index similar to that observed in our study. The method employed here by us has the advantage of simplicity as well as controlled and reproducible spheroid size. Reproducible size is an important factor in standardisation as large spheroids may undergo core necrosis (53), whereas variation in size may lead to differences in cell-cell coupling (54, 55) and insulin secretion (56). An attractive alternative, especially when using extracellular electrophysiology, may be given by cell electrophoresis of dispersed cells onto electrodes (15).

Clearly spheroid formation provided far more robust electrophysiological responses and even permitted to reliably detect action potentials which are difficult to monitor in monolayers even when using electrodes coated with a conducting polymer (15). In all EndoC-βH models and in primary islet cells we observed a decrease in SP frequency and an increase in their amplitude. The changes in amplitudes of SPs in 3D versus 2D cultures are most likely due to improved coupling (11) and correlated with an increase in insulin secretion in the three cell models tested. The improvement in insulin secretion observed here was comparable to published data (6, 28). Notably, in EndoC-βH5 cells a biphasic pattern of electrical activity was apparent upon glucose stimulation, which is a hallmark of primary islets (14) and such a pattern had also been reported in dynamic insulin secretion assays of EndoC-βH5 cells (9). We did not observe any significant effect on slow potentials by activating the GLP-1 receptor via its native agonist at physiological concentrations. Note that we previously observed that these concentrations significantly increase slow potential amplitude and frequency in mouse and human islets although the effect was small (14). The effects of GLP-1 on action potentials observed here indicate that the hormone signalling pathway was active in the clonal EndoC-βH cells in our study. As the GLP-1 inactivating protease DDP-4 is expressed in EndoC-βH cells (57), most studies used the incretin mimetic peptide exendin-4 (6, 9, 28, 58) with one exception (59). However, we think that such a pharmacological approach yields less insight. In the same vein, glucose concentrations should be chosen carefully and we prefer a minimal glucose concentration of 3 mM to its complete absence. Although stimulation indices are more pronounced when referring to 0 glucose, prolonged absence of glucose may considerably alter gene expression profiles (60).

We were not able to form stable spheroids using rat insulinoma INS-1 cells (48), a widely used and relevant clonal β-cell model. Most spheroids published of this cell line were generally of variable sizes and poorly defined borders although the methods used for spheroid formation provided useful spheroids in another β-cell lines such as MIN6 (30, 61-63). A more recent publication reported the generation of uniform spheroids but were maintained in a device that is not compatible with electrophysiological recordings (64, 65). It is also of note that most of the previously reported INS-1 aggregates showed either a considerable right shift of glucose dependency or a stark reduction in glucose induced insulin secretion (61, 62). Cx36 overexpression induced here a coherent pattern in electrical activity and insulin secretion: an increase in glucose-sensitive cells, reduced basal electrical and secretory activity and consequently almost doubling in glucose induced stimulation indices. These observations are in line with the general function of Cx36 in islets (18) and in INS-1(66).

In conclusion, EndoC-βH1 and especially EndoC-βH5 spheroids provide a very useful model to test the effect of different variables on electrical activity and physiological cell-cell coupling by MEA analysis in line with the advocated utility in drug testing (6). Cx36-transduced INS-1 cells may be suitable, but obviously restricted to rodents. These models may also be of interest as biological substrate for organs on chips and microorgan-based sensors for continuous nutrient sensing (36, 67, 68).

## Author Contributions

The study was designed and funding was obtained by JL; EP, KLF, JG, ML, PAS, MR and JL researched and analysed the data; JL, EP, KLF, JG, ML, PAS, MR cowrote the manuscript; All authors have consented to the submitted manuscript. EP and KLF; equal contributions.

## Acknowledgments

We thank the vectorology platform Vect’UB (CNRS UMS3427, INSERM US005, Univ. Bordeaux) for providing lentiviral particles and technical support. We thank Thierry Leste Lasserre (Univ. Bordeaux, INSERM, PUMA, Neurocentre Magendie, U1215) for help with the qPCR. We are grateful to Human Cell Design (Toulouse, France) for providing us with EndoC-βH1 cells and OPTIβ1 medium.

## Data Availability Statement

Data will be made available upon reasonable request.

