## Supplemental Material for "Extracellular electrophysiology on clonal human β-cell spheroids"

**ELECTRONIC SUPPLEMENTARY MATERIAL**

**SUPPL FIG 1**

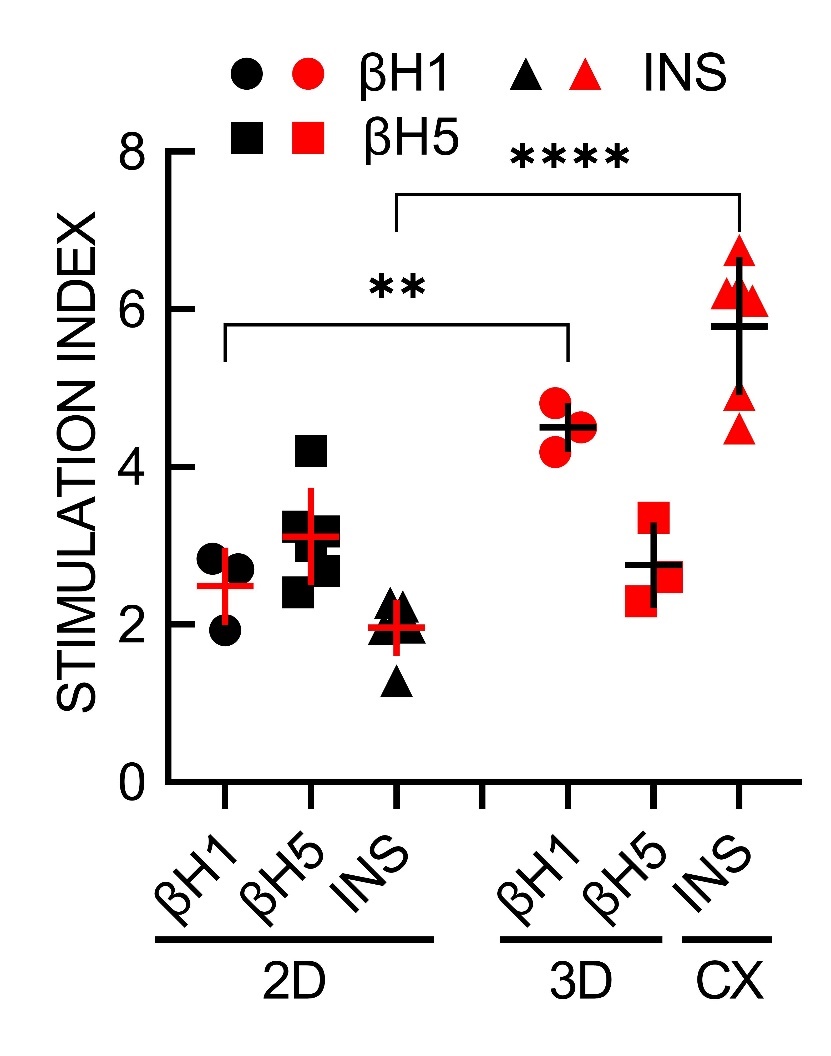

**Supplemental Figure 1:** Relative stimulation indices. EndoC-βH1 and EndoC-βH5 fold stimulation of 11 mM vs 3 mM glucose, for INS-1 fold stimulation of 15 mM vs 3 mM glucose. Black symbols, 2D monolayer; red symbols, 3D spheroids or CX overexpression; CX, connexin36 overexpression. Tukey post-hoc test; **, 2p< 0.01; ****, 2p< 0.0001; n 3 to 6.

**SUPPL. FIG 2**

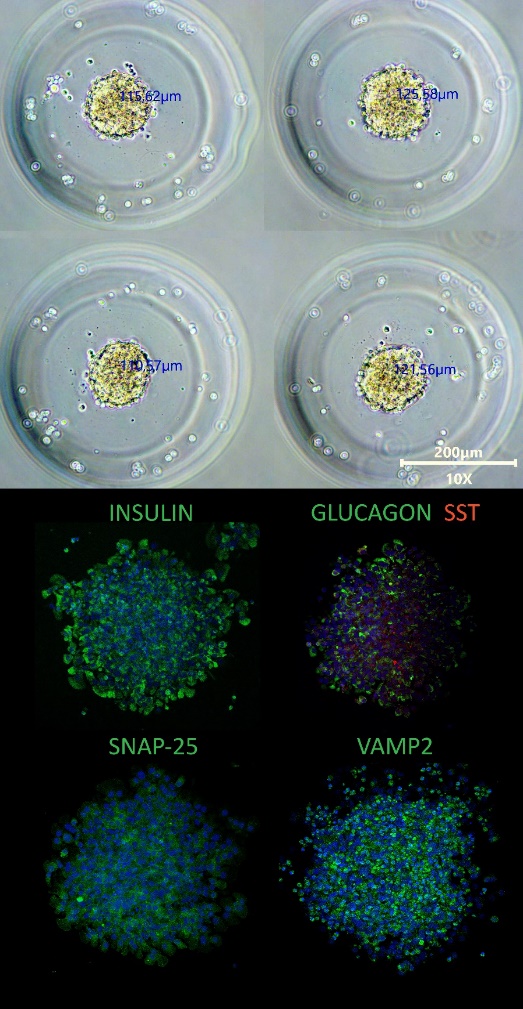

**Supplemental Figure 2:** EndoC-βH5 spheroids. Upper panel: representative images of EndoC--βH5 spheroids; lower panel, immunostaining for insulin, glucagon/somatostatin, SNAP-25 or VAMP2. Nuclei were stained with DAPI (blue).

**SUPPL FIG 3**

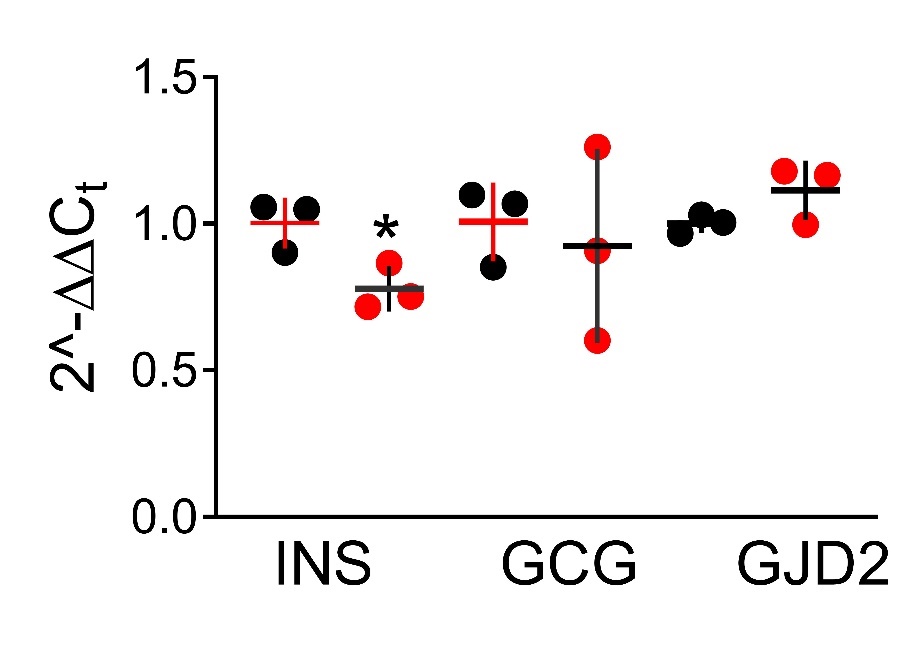

**Supplemental Figure 3:** Relative changes in expression values for preproinsulin (INS), preproglucagon (GCG) or connexin 36 (GJD2) in EndoC-βH5 monolayers (black circles) or spheroids (red circles). Mean C_t_ values in spheroids were (±SD; n=3): *YWHAZ*, 22.39±0.17; *GAPDH* 20.32±0.17; *INS* 16.67±0.22; *GCG* 27.51±0.23; *GJD2* 25.90±0.17; *FAP* and *IRX2* did not provide any amplification in spheroids or monolayers.

Table 1. qPCR primer sequences

| **Gene** | **GenBank ID** | **Forward sequence (5'-3')** | **Reverse sequence (5'-3')** |
| --- | --- | --- | --- |
| *YWHAZ* | NM_003406 | CCTGTTTTAGCCTTCTGTCTTGTCA | ATGCGGCCTTTTTCCAAGTAC |
| *GAPDH* | NM_002046 | CACCCATGGCAAATTCC | TGGGATTTCCATTGATGACAAG |
| *INS* | NM_000207 | TGGAGGGGTCCCTGCAGAA | AGTAGTTCTCCAGCTGGTAGA |
| *GCG* | NM_002054 | ATCTTCACAACATCACCTGCTA | TACACCTCTTAAATTTACAGGACTT |
| *GJD2* | NM_020660 | CAGCTTGTGGACTTTGGTT | GTGGCTCAGCTAGACACTCT |

### 1 Samples were homogenized in Tri-reagent (Euromedex, France). RNA was isolated using a standard chloroform/isopropanol protocol (1) and purified by incubation with Turbo DNA-free (Fisher Scientific). RNA was processed and analyzed following an adaptation of published methods (2). cDNA was synthesized from 0.5 μg of total RNA using Maxima Reverse Transcriptase (Fisher Scientific) and primed with oligo-dT primers (Fisher Scientific) and random primers (Fisher Scientific). qPCR was perfomed using a LightCycler® 480 Real-Time PCR System (Roche, Meylan, France). qPCR reactions were done in duplicate for each sample, using transcript-specific primers, cDNA (4 ng) and LightCycler 480 SYBR Green I Master (Roche) in a final volume of 10 μl. Primers sequences are reported in table 1. For the determination of the reference gene, the refFinder method was used (3). Relative expression analysis was normalized against two reference genes: tyrosine 3-monooxygenase/tryptophan 5-monooxygenase activation protein zeta (Ywhaz) and glyceraldehyde-3-phosphate dehydrogenase (Gapdh). The relative level of expression was calculated using the comparative 2-ΔΔCT method (4).

1. Chomczynski P, Sacchi N. Single-step method of RNA isolation by acid guanidinium thiocyanate-phenol-chloroform extraction. Anal Biochem. 1987; 162:156-9

### 2. Bustin SA, Benes V, Garson JA, Hellemans J, Huggett J, Kubista M, Mueller R, Nolan T, Pfaffl MW, Shipley GL, Vandesompele J, Wittwer CT. The MIQE guidelines: minimum information for publication of quantitative real-time PCR experiments. Clin Chem. 2009; 55: 611-22.

### 3. [F Xie, P Xiao, D Chen, L Xu, B Zhang. miRDeepFinder: a miRNA analysis tool for deep sequencing of plant small RNAs. Plant Mol Biol. 2012 ; 80 : 75-84.](http://www.ncbi.nlm.nih.gov/pubmed/22290409)

4. Livak KJ, Schmittgen TD. Analysis of relative gene expression data using real-time quantitative PCR and the 2(-Delta Delta C(T)) Method. Methods. 2001; 25:402-8.
